# Which food patches are worth exploring? Foraging desert birds do not follow environmental indicators of seed abundance at small scales: a field experiment

**DOI:** 10.1101/295923

**Authors:** Fernando A. Milesi, Javier Lopez de Casenave, Víctor R. Cueto

**Affiliations:** Desert Community Ecology Research Team (Ecodes), Departamento de Ecología, Genética y Evolución, Facultad de Cs. Exactas y Naturales, Universidad de Buenos Aires, and IEGEBA (UBA– CONICET), Buenos Aires, Argentina; Present address: Instituto de Investigaciones en Biodiversidad y Medio Ambiente, INIBIOMA (UNCo– CONICET), Centro de Ecología Aplicada del Neuquén (CEAN), Junín de los Andes, Neuquén, Argentina; Present address: Centro de Investigación Esquel de Montaña y Estepa Patagónica, CIEMEP (UNPSJB– CONICET), Esquel, Chubut, Argentina

## Abstract

Consumers should show strong spatial preferences when foraging in environments where food availability is highly heterogeneous and predictable from its correlation with informative environmental features. This is the case for postdispersal granivores in most arid areas, where soil seed bank abundance and composition associates persistently with vegetation structure at small scales (e.g., decimeters to meters). We analysed seasonal single-seed removal by granivorous birds from 300 experimental devices in the algarrobal of the central Monte desert. Spatial selectivity was analysed by comparing the structural characteristics of used vs. available microhabitats and evaluated against bottom-up and top-down hypotheses based on our previous knowledge on local seed bank abundance, composition and dynamics. Seed removal, which showed its expected seasonal variability, was also explored for spatial autocorrelation and environmental dependencies at bigger scales. Postdispersal granivorous birds were less selective in their use of foraging space than expected if patch appearance were providing them useful information to guide their search for profitable foraging patches. No kind of microhabitat, as defined by their vegetation and soil structure, was safe from bird exploration. The only consistent selective pattern at this scale was closer to a top-down spatial effect by birds, i.e., a cause (and not a consequence) of the seed bank dynamics. Bigger spatial scales proved more relevant to describe heterogeneity in the use of foraging patches in this habitat. Closeness to tall trees, probably related to bird territoriality and reproduction or to their perception of predation risk, seems to determine a first level of selection that defines explorable space, and then microhabitat openness exerts an influence on which patches are effectively exploited (or more frequently explored) among those accessible.

## INTRODUCTION

The decision-making process of foraging animals involves gathering information on critical factors, either by inference from reliable environmental cues or by local assessment after a site has been explored (Mitchell 1989, Valone 1991, Stephens 2007). Food availability, together with costs and benefits related to foraging efficiency, vulnerability to predators and microclimate are usual candidate factors affecting selection of foraging patches by small individual animals (Wiens 1985, Repasky and Schluter 1994, Meyer and Valone 1999). The ability to assess and respond to patchiness at small scales allows foragers to exploit their habitat more efficiently, particularly if there are physical cues associated with patch boundaries that can be perceived through direct sensory input (Bell 1990, Schmidt & Brown 1996, Fierer & Kotler 2000). Consumers should show strong spatial preferences when foraging in environments where food availability is highly heterogeneous and predictable from its correlation with informative environmental features. This would allow them to behave as “prescient foragers”, releasing them from the costs of random or indiscriminate exploration (paying ‘the penalty of ignorance’) or continuously gathering spatial data to update their knowledge on distribution and location of food patches (Valone 1991, Kohlmann and Risenhoover 1998, Dall et al. 2005, Olsson & Brown 2006, Eliassen et al. 2007, Olsson & Molokwu 2007, Sih 2011, Marshall et al. 2013).

Postdispersal seed predation, as other foraging situations, can be split in two components: (1) patch exploration, stated as the probability of at least one seed being removed from a patch, and (2) patch exploitation, the amount or proportion of seeds removed once the patch was explored (‘seed encounter’ and ‘seed exploitation’ *sensu* Hulme 1994). Seed availability in the soil of most deserts is highly heterogeneous at small scales (e.g., decimeters to meters) and associates persistently with vegetation structure (e.g., Nelson & Chew 1977, Price and Reichman 1987, Guo et al. 1998, Caballero et al. 2008), and the central Monte Desert (Argentina) is no exception: seeds and litter consistently accumulate under shrub and tree canopies (Marone & Horno 1997, Marone et al. 2004, Milesi & Marone 2015, Andrade 2016). Woody cover (and litter) provides conspicuous environmental cues of resource abundance that should allow visual foragers such as birds to increase their foraging success by exploiting this “pre-harvest” or “prior” information accumulating their previous experiences and evolutionary history (Valone 1991, Dall et al. 2005, Marshall et al. 2013). As predicted by classic ‘attack’ foraging models like the ‘optimal patch choice’ and ‘diet’ models (Pyke 1984, Stephens and Krebs 1986, Ydenberg et al. 2007), postdispersal granivorous birds should follow patch appearance and neglect “poor patches” before going through the cost of their exploration. However, the expected association between vegetation cover and postdispersal seed consumption was not evident at this spatial scale when tested by following foraging granivorous birds at field: no differences in vegetation characteristics were found between foraging microsites and randomly-chosen microsites (Milesi et al. 2008).

Foraging decisions informed by environmental cues of resource abundance may have further ecological consequences (Schmidt et al. 2010) if selective consumption influences the spatial distribution of resources (i.e., a top-down spatial effect). Non-random seed removal affects the composition and heterogeneity of the seed bank, so spatial patterns of seed abundance on the ground and, later, of adult plants may be interpreted also as the consequence —instead of the cause— of consumption by granivores (Hulme 1997, Russell and Schupp 1998, Guo et al. 1995, Howe & Brown 2001, García et al. 2005, Orrock et al. 2006, Maron et al. 2012). Granivorous birds should be able to modify abundance, composition and spatio-temporal heterogeneity of the soil seed bank in arid areas where they are important granivores (e.g., at the Monte desert: Marone et al. 1998b, 2008), especially because their diet is usually selective (e.g., Willson 1971, Pulliam 1985, Cueto et al. 2006, Camin et al. 2015). However, most studies on the impact of granivores on desert seed banks have emphasised the impact of rodents and ants (e.g., Nelson & Chew 1977, Reichman 1979, MacMahon et al. 2000, Anderson & MacMahon 2001), mostly as a derivation of the presumed “irrelevance” of birds on deserts of the Northern Hemisphere (Mares & Rosenzweig 1978, Abramsky 1983, Parmenter et al. 1984; but see Thompson et al. 1991, Guo et al. 1995, Lopez de Casenave et al. 1998, Marone et al. 2000b).

We evaluated spatial selectivity at small scales by foraging postdispersal granivorous birds in open woodlands (*algarrobales*) of the central Monte desert. Every season, we analysed the spatial pattern of seed removal from experimental devices and compared the structural characteristics of the vegetation in used microhabitats against those available. Most studies on seed removal emphasize the exploitation component of seed predation through explicit or implicit baiting (e.g., proper baiting, training sessions, ad-libitum patches) to minimize the probability that focal organisms do not detect or disregard the experimental setup. Instead, we offered a single seed per device aiming to detect where do birds normally search for seeds, and not where or how much they are able to remove when an extraordinary rich patch is suddenly available (the “magic pudding effect”: Bedoya-Pérez et al. 2013). Though not frequently, some studies have used, distinguished or compared both approaches (e.g., Thompson et al. 1991, Manson & Stiles 1998, Kelt et al. 2004). According to our previous knowledge on local seed-bank dynamics, (Marone & Horno 1997, Marone et al. 2004, Milesi 2006) the bottom-up hypothesis predicts that birds should remove experimental seeds preferentially from devices under shrubs and trees or under grass cover (i.e., where seeds concentrate). The top-down interpretation of differences between potential and effective seed banks (Marone et al. 1998a, 1998b, 2000a) calls instead for granivorous birds that use open, bare-soil microhabitats disproportionally, at least during autumn and winter, to account for the observed postdispersal reduction in the density of their preferred seeds in open microsites. Use of space at small scales may not only vary with time but depend on context at bigger spatial scales, a key knowledge for proper interpretation of selective patterns (Kotliar & Wiens 1990, Jones 2001, Mayor et al. 2009). Seed removal from our experimental devices was explored for spatial patterns and environmental dependencies, particularly because we had previous observational evidence on the relevance of tall trees on granivorous birds foraging in this habitat (Milesi et al. 2008).

## METHODS

### Study area

The study was done in the Biosphere Reserve of Ñacuñán (34°03’S, 67°54.5’W), in the central Monte desert, Province of Mendoza, Argentina. The climate is dry, with wide variations in annual precipitation between years (mean: 348.9 mm, range: 192.6–585.4 mm, 1972–2002). It is also highly seasonal, with warm and rainy summers (>20 °C, 269 mm) and cold and dry winters (< 10 °C, 80 mm). A complete description of the area can be found in Lopez de Casenave (2001).

The main habitat of the Reserve is the *algarrobal*, an open woodland of algarrobo (*Prosopis flexuosa*) trees 3–6 m high scattered in a matrix of perennial *Larrea divaricata* tall shrubs (1–3 m high, horizontal cover >35%). Other woody species are *Geoffroea decorticans* trees, tall shrubs such as *Capparis atamisquea*, *Condalia microphylla* and *Atriplex lampa* (usually >1 m high), and low shrubs (~20% cover, usually <1 m high) such as *Lycium* spp., *Verbena aspera* and *Acantholippia seriphioides*. There is also an important cover (>25%) of perennial grasses (*Pappophorum* spp., *Trichloris crinita*, *Digitaria californica*, *Aristida* spp., *Setaria* spp., *Sporobolus cryptandrus*). Most of the reserve has been closed to cattle ranching and other significant human activities since 1971. About a third of the surface of the algarrobal lacks perennial vegetation in the form of open patches of variable size (from centimeters to meters). Forb cover is highly variable among seasons and years, usually an order of magnitude lower than grass cover; therefore they were not considered in the description of the vegetation structure, following local studies of the seed bank (Marone & Horno 1997, Marone et al. 2004) and bird foraging (Milesi et al. 2008).

Seeds of herbaceous plants are the staple diet of granivorous birds (75–99% of their granivorous diet is grasses and one forb species; Marone et al. 2008). Their abundance in the soil is very heterogeneous at small scales, with close patches of extreme abundances. Seeds are consistently more abundant under trees and shrubs and in depressions of the soil where litter accumulates, mostly because of the persistent seed bank of forb seeds (Marone et al. 2004, Andrade 2016). The abundance of grass seeds, preferred by these birds (Cueto et al. 2006, Camin et al. 2015), is less heterogeneous though still higher under woody cover, with some intra- and inter-annual variability. Forbs start producing seeds in the spring and grasses usually in summer. Primary seed dispersal starts in late spring and finishes by winter; maximum seed availability in the soil occurs during autumn and winter, with a minimum when summer begins (Marone et al. 1998a, 2004).

### Experimental offer of single seeds

Single seeds were offered on top of each of 300 devices, made of upside-down feet of plastic flute glasses with their stems buried so the top (originally the base of the glass) remained 2–3 cm over the ground. This configuration prevented access by granivorous ants and other arthropods that cannot walk upside-down on the smooth plastic surface (Kelrick et al. 1986, Lopez de Casenave et al. 1998, *pers. obs.*). Devices were arranged 5-m apart on three 10×10 grids (‘J’, ‘F’ and ‘V’; area ≈ 2000 m^2^ each) located 80–400 m apart within the algarrobal. A single *Setaria italica* seed (a commercial species bigger than the otherwise similar local *Setaria leucopila*, both readily consumed by birds; Cueto et al. 2006) was offered on each device for two consecutive days from sunrise to sunset (standardised following the civil twilight data and criteria by U.S. Naval Observatory; U.S. Navy n.d.), once per season (i.e., ~23.3 h in Autumn to ~29.5 h in Summer). The top surface of the device, 6 cm in diameter, was covered with local soil for a similar visual appearance to the surrounding ground. According to estimations from simultaneous soil samples (L. Marone, unpublished), our experimental seed offer should not result particularly attractive against background seed offer: from 2.4 (bare soil in spring) to 19.3 (beneath shrubs in winter) grass seeds are expected in a similar sized area of soil. The experimental single seed on the device surface had a similar biomass density to the expected average for grass seeds in this habitat during winter (~0.1 mg/cm^2^) and 36% of the biomass of all consumable seeds (those in the diet of granivorous birds, Marone et al. 2008).

Rodents that may have remove seeds in the area are mainly nocturnal (Lopez de Casenave et al. 1998). Two additional trials were done under a modified protocol to test the assumption that birds were the (only) diurnal organisms removing seeds from these devices. Clayish local soil was smoothed around 50 devices in grid F before offering seeds during an extra day after the main summer and winter trials. Footprints of birds, mammals and other taxa (insects and lizards) were easily recognised on the smooth fine soil. In most cases where the seed has been removed only bird footprints were detected, in both winter (32/35 = 91%) and summer (7/9 = 78%) trials. The rest had mixed footprints of birds and other taxa or not recognisable footprints, but no device without seed had footprints of other taxa exclusively. On the other side, no bird footprints were found around devices where the seed had remain, suggesting that the seed was usually removed when closely approached by walking birds.

Seed removal from these devices can be assigned to small seed-eating birds of the ground foragers guild, particularly *Zonotrichia capensis*, *Saltatricula multicolor*, *Diuca diuca*, *Poospiza ornata* (in summer) and *Phrygilus carbonarius* (Marone 1992, Lopez de Casenave et al. 2008, Milesi et al. 2008). Other birds in Ñacuñán do not for forage for seeds on the ground or are rare or occasional visitors to this habitat (Lopez de Casenave 2001; e.g., *Poospiza torquata*, *Catamenia analis*, *Zenaida auriculata*, *Columba maculosa*, *Columbina picui*, *Carduelis magellanica*, *Molothrus bonariensis*, *Passer domesticus*, *Eudromia elegans*, *Nothura darwinii*). The expectation for these other species to prefer microhabitats with more seeds is still valid, so any occasional bird removing seeds from the devices should not alter our main conclusions.

Since seed removal from a device was not independent between the two consecutive days in any of the four seasons (Fisher exact tests for 2×2 contingency tables: χ^2^ > 14.15, *P* < 0.001, *n* = 300, with more observations of “double removal” and “never removed” than expected by chance), a device was defined as “used” in each season if the seed was removed at least once during the two days it was offered. Independence of seed removal among seasons was tested for each grid by comparing the distribution of observed frequencies of the number of season in which each device was used (0–4) against the expected frequencies calculated as the product of four Bernoulli trials with *n* = 100 (each seasonal experiment) with variable probabilities of success, estimated as the proportion of used devices in that grid for each season. Goodness of fit was evaluated with a χ^2^ statistic.

Temporal and spatial heterogeneity in intensity of seed removal (proportion of used devices per grid) was tested with binomial generalised linear models (*logit* link) with GRID and SEASON as independent categorical variables. Significance of predictors was assessed comparing the change in deviance of nested models obtained through stepwise backwards elimination, asymptotically distributed as χ^2^. The ratio of deviance to degrees of freedom in the minimum adequate model was 1.74, but corrections for overdispersion did not change interpretation of results (*F*- vs. χ^2^-tests).

### Vegetation and soil characteristics at the microhabitat scale

Studies on use of space through short-term observations rely on asymmetric evidence: while patch use can be inferred from seed removal, non-removal does not imply the patch is not to be explored eventually. This lack of a proper “no use” group should raise concerns on simple statistical analyses based on a priori classification into exclusive groups (e.g., discriminant analyses), particularly if they assume similar variability in both (homocedasticity), or based on assigning zero probability of use (e.g., classical logistic regression). Identification of explanatory variables and predictive value of the statistical models can both suffer from the unrecognized probability of false-negatives (Tyre et al. 2003, Martin et al. 2005, Elith et al. 2006). Though more complex modelling strategies can incorporate or simulate incomplete evidence on absences in a spatially explicit context (e.g., species distribution and Bayesian models; see Hirzel et al. 2002, Elith et al. 2006, Latimer et al. 2006), we preferred an indirect strategy to evaluate selection that best matches how we developed our set of hypotheses and predictions. We started by detecting and simplifying the main structural and floristic characteristics defining the heterogeneity of this habitat at the microsite scale, which we assumed in previous studies (e.g., Marone et al. 2004, 2008, Milesi et al. 2008) to depend mainly on shrub and tree cover and relate with main characteristics of the seed bank. Then, we evaluated if used microsites (i.e., those where the seed had been removed) were a random (no selection) or a skewed (selective) sample of available microsites, both on their characteristics and on their spatial position.

Characteristics of the vegetation and soil cover in the microhabitat around each device position were measured by placing a vertical 1-m long pole (2 cm diameter) every 10 cm along four 50-cm transects (=20 points per microsite) from the device to each cardinal direction. At each point, perennial plants (trees, shrubs and grasses) touching the pole were identified to genus level. The presence of vegetation >1 m and the presence of dense litter (when it prevented from seeing the mineral soil below) or its absence (bare soil) were also recorded. Percentage cover per plant group (grasses, standing dry grasses, low shrubs, tall shrubs, and trees) and of bare soil and deep litter were calculated after those measurements.

A Principal Components Analysis (PCA) with Varimax rotation of the selected axes was done to reduce the number of dimensions of the ten variables measured at the microsite scale. Some variables were previously transformed (arcsin, square root or logarithm) to improve the symmetry of their distributions and then standardised into the PCA correlation matrix.

Alternative analyses at the level of plant genus gave noisier but similar results on the main axes. Three components (PC1–PC3) were retained following the Kaiser criterion (eigenvalue >1), the broken-stick model, and the scree-plot (Jackson 1993). Before multidimensional analyses, scores on each axis were multiplied by its eigenvalue to weight them according to the variability they represent (see Peres-Neto & Jackson 2001); though applied for correctness, these variable transformations and weights did not change results significantly. Separate PCA analyses for each grid resulted in similar principal components and scores correlated with those of the grouped analysis (Pearson correlation, all cases *n* = 100, *r* > 0.8, *P* < 0.001, except for PC3 in grid V: *r* = 0.21, *P* = 0.03) confirming that heterogeneity of main characteristics at this scale were similar for the three sites. Therefore, only PCA scores based on all microhabitats were used for subsequent analyses. Scores based on microhabitat characteristics were compared among grids with Kruskal-Wallis tests.

The microhabitat around each device was also categorized at field *a priori* following the same criteria used on previous studies (Marone & Horno 1997, Marone et al. 2004): beneath trees, beneath tall shrubs, beneath low shrubs, beneath grasses (under no woody cover), and bare soil (no perennial cover). Characteristics of the vegetation around 79 grid points (26%) did not fit neatly into any of those categories (e.g., shrub borders); they were assigned to an “intermediate” category.

Big trees (trees >3 m and algarrobos >4 m high) in and around the grids were mapped, measuring the distance between each device and the nearest tree canopy. Other minimum distances to vegetation in several height strata and to closest canopy of each plant group were measured but resulted highly correlated with measurements of horizontal cover (since most distances were smaller than a microsite radius, e.g., 96% of distances to nearest grass or 78% to vegetation 1–2 m high, see Milesi 2006). Therefore, only distances to tall trees were informative in addition to the measured characteristics at the microhabitat scale. Distances were compared among grids with Kruskal-Wallis tests.

### Foraging site selection based on microhabitat characteristics

Selection at the microhabitat scale was evaluated (1) multidimensionally, with a spatial technique applied to results of the PCA ordination and (2) unidimensionally, with a randomization test for each of the three retained PCA components. The first test is analogue to the representation of used and available microsites in multidimensional scatterplots and the evaluation of the spatial segregation between two classes of points. The test was a 3-D extension of a 2-D point pattern spatial analysis that classifies each point by its type and that of its nearest neighbour and compares the proportion of each kind of pair with that expected by chance (i.e., a join-counts analysis of a binary label according to a nearest-neighbour matrix, testing for differences against a random labelling model: Dale et al. 2002, Fortin & Dale 2005). The number of pairs of equally labelled points will be greater than expected by chance if used points are aggregated or if there are available zones where there is no use (i.e., available points are aggregated). Global spatial segregation between classes of points was evaluated with a 2×2 contingency table, with expected frequencies and statistic (*C*, asymptotically distributed as chi-square with 2 d.f.) as proposed by Dixon (NN test; Dixon 1994). When significant evidence of global segregation was found, each type of pair was tested with an asymptotically normal *Z* statistic (Dixon 1994). Statistical analysis were done in R (R Core Team 2013), modifying the functions provided by P. Dixon (Dixon n.d.) to a multidimensional case to obtain the matrix of Euclidian distances between points and identify nearest neighbours in a 3-D case.

To evaluate selection on each principal component, the null hypothesis that microhabitat characteristics of used devices are a random sample of those of the available ones was tested with randomization tests (Crowley 1992, Potvin & Roff 1993, Adams & Anthony 1996). Statistics of central tendency (mean, median) and dispersion (variance) were calculated from 4999 or 1999 samples, respectively, of the same size as observed (used), taken without replacement from the available values, to evaluate selection consisting in a skewed use of lower or higher values of the environmental variable (resulting in lower or higher mean or median) and selection consisting in avoiding extreme or central values (lower or higher dispersion, respectively; see James & McCulloch 1990, Clark & Shutler 1999, Hirzel et al. 2002). Results based on the median of the distributions were very similar to those based on the mean, so they are not presented. A pseudo-*P* value associated to the hypothesis that the observed statistic was obtained by chance was calculated as double the number of equal or more extreme values than the observed in the distribution, divided by the number of samples taken including the observed (i.e., a two-tailed test). Spatial autocorrelation of PCA scores was rare (see below), so no correction was applied to statistically evaluate selection hypotheses at the microhabitat scale.

Two sets of confidence intervals and probabilities were calculated, based on different null models. First, randomly chosen values were obtained in each iteration from the 300 values of all available microhabitats to evaluate selection assuming no selective use of space at bigger scales (up to the extent of the study). Second, a stratified null model was done to control for a possible habitat selection at the grid scale (i.e., assuming a hierarchical use of space based on the observation that grids differed systematically in the proportion of used devices, see below). Under this model, random samples of the same size as observed were taken from each grid, so the expected mean value was an average of the mean values of the three grids weighted for the number of used devices in each of them.

### Foraging site selection based on device position

To analyse the influence of location (cf. microhabitat features) on the use of a device, seed removal in each 10×10 grid was examined for spatial autocorrelation using spatial analyses for non-continuous data, assuming an isotropic process. Spatial autocorrelation of seasonal seed removal and of microhabitat characteristics were evaluated comparing a measure of similarity between pairs of points given by their position with another determined by the focal variable (agreement between two matrices of similarity; Dale et al. 2002). Discrete Euclidian distances between devices (from regular grids) were aggregated in four distance classes up to half the size of a grid side: <8.5 m (nearest 8 neighbours of a focal central grid point), 8.5–12.5 m (12 neighbours), 12.5–17 m (16), and 17–22 m (24). Relationship between points given by distance (known as matrix of weights, *W*) were considered binary, with 1 indicating that two points are separated by a distance in the focal distance class, and 0 otherwise. Two similar spatial analyses were used according to the type of variable (Dale et al. 2002, Fortin et al. 2002, Perry et al. 2002). An analysis of join-counts was used to evaluate spatial autocorrelation of seasonal seed removal (a binary variable) in each distance class (Rosenberg 2001). This is simply counting the number of points joined (i.e. they are separated by a distance in the focal distance class) that share a property (i.e., both have been used in the same season). For continuous variables (values of the main PCA axes, distance to trees), Moran’s *I* statistic was used. Results from both analyses are shown in correlograms which can be interpreted in a similar way: values higher than expected indicate a positive spatial autocorrelation at the focal distance class (i.e., aggregated pattern), and values significantly low indicate negative spatial autocorrelation (i.e., overdispersed or regular pattern).

The number of pairs of devices that were both used (1–1 joins) was compared against expectations from two different models. The first one is a model of complete spatial randomness (CSR) in which the probability of use of a device follows a homogeneous Poisson process depending only on the observed number of devices used in a grid, with no spatial interaction (i.e., all devices have the same chance of being used independently of its neighbours). The second model (distance-to-tree model, DTT) is a very simple heterogeneous Poisson process to illustrate aggregation of seed removal from first order or induced autocorrelation associated with distance to tall trees. This analysis tests the observed configuration keeping the observed composition, edge effects and potential habitat selection at bigger scales (i.e. different use of grids), assuming no second order autocorrelation (i.e., the use of a device is independent of that of its neighbours except for the modelled first order autocorrelation). The expected distribution of the statistic under each model was estimated with 1999 random samples of grid points of the same size as observed (with no replacement). For CSR model, all points shared a single probability to be selected. For DTT model, probability of use was chosen to vary discretely and inversely with distance to the nearest tall tree: a device had a relative probability of use of 0.6 if at <5 m or 0.3 if at 5–10 m of an algarrobo >4m high and a probability of 0.1 if at <10 m to the nearest tree >3 m high; points at >10 m of any tall tree (none in grid J, 4 in F and 26 in V) had no chance of being selected (probability = 0). Expected values (median) and limits of confidence intervals (percentiles 2.5% and 97.5%) were obtained from each of the generated distributions, estimating the probability *P* that the observed value belongs to the distribution under the expected model as two times the proportion of equal or more extreme values than the one observed, including it (i.e., a two-tailed test with *n* = 2000). Algorithms for resampling and join-counts estimation were programmed in R (R Core Team 2013). Moran’s *I* statistics for continuous variables were analysed in a similar way, comparing them against a CSR model as implemented in the software PASSAGE (Rosenberg 2001). Complementary spatial analyses with alternate techniques (SADIE: Perry & Dixon 2002) provided similar evidence (see Milesi 2006).

## RESULTS

The number of devices where the single seed was removed varied among seasons and grids. Seed removal was higher in autumn and winter and decreased in spring to summer in each of the three grids, which showed a consistent relative level of use: J had always higher seed removal, followed by F, and then V (Fig. 1). Both SEASON and GRID (and not their interaction) were relevant predictors of the number of devices used, with the spatial position at GRID scale an even stronger predictor than SEASON (Table 1).

**Table 1.**
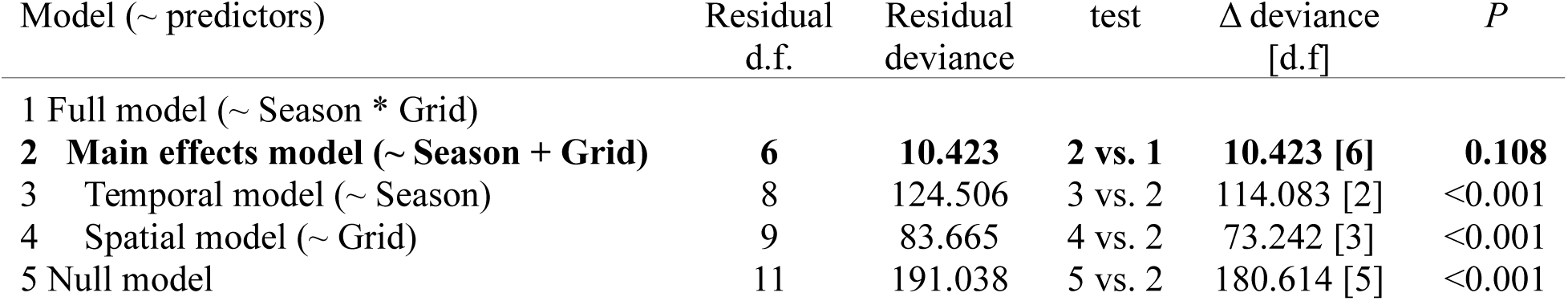
Analysis of deviance of nested binomial GLMs (*logit* link) obtained through backwards elimination from the full model, with temporal (SEASON, four categories) and spatial (GRID, three categories) heterogeneity as predictors of the proportion of used devices in each experiment. Minimum adequate model is shown in bold and includes both main effects (model 2).

**Figure 1.**
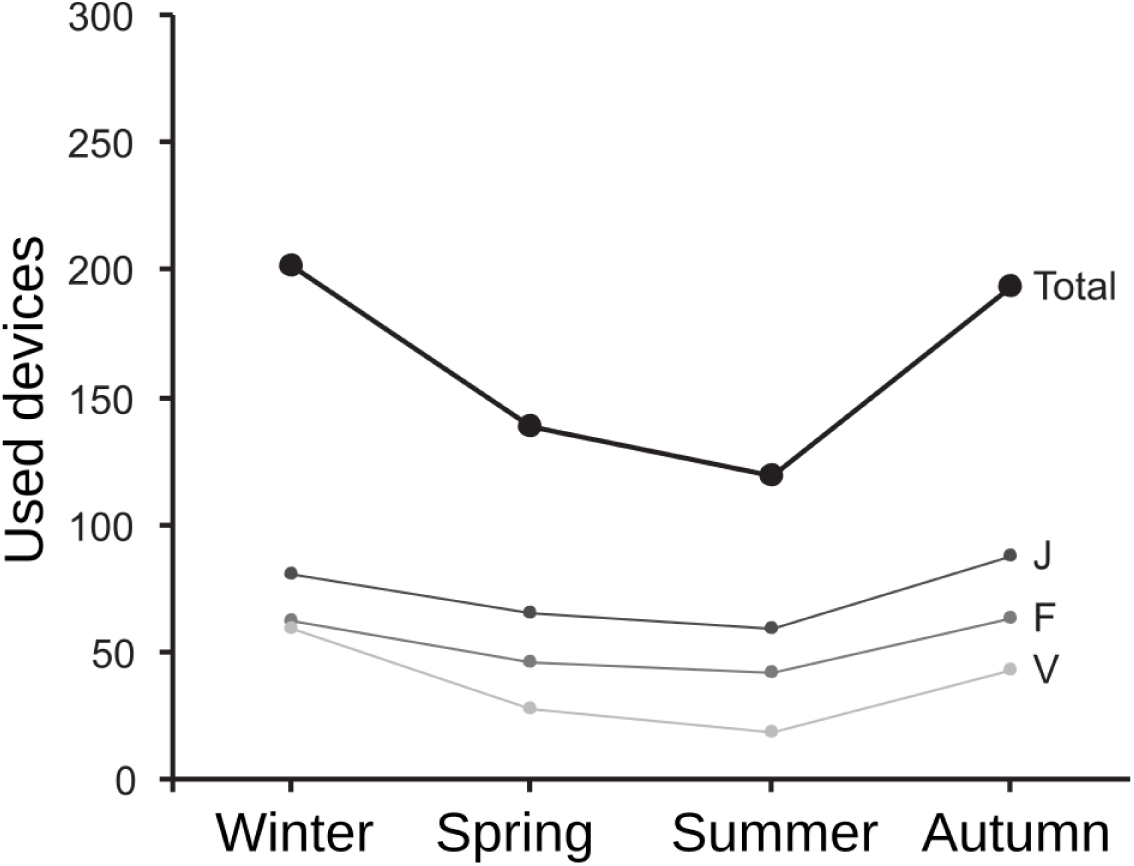
Number of devices where the seed was removed in at least one of the two days offered per season and grid. Total number of devices available was 300 in each season, arranged in three 10×10 grids (J, F, V).

Seed removal from a particular device in a grid was not independent among seasonal trials (J: χ^2^_4_ = 80.23, F: χ^2^_4_ = 19.90, V: χ^2^_4_ = 42.02; all: *n* = 100, *P* < 0.001). There were more observations in the categories “always used (4/4)” and “never used (0/4)” than expected if seed-offer devices in a grid were randomly and independently used each season.

The first three components of PCA on ten characteristics of vegetation and litter measured at the microhabitat scale (around seed offer devices) retained 73% of the variability in the correlation matrix (Table 2). The first component (PC1) represented general vegetation cover, with positive values associated with tall shrubs and litter and negative values with open areas and bare ground. The second (PC2) was associated with tree cover (to the positive values), and the third (PC3) with the rest of perennial vegetation: grasses to the positive and low shrubs to the negative values. These components can be associated with the soil seed bank according to previous local results based on sampling microhabitats categorized *a-priori* at field: total seed abundance in the soil should be associated with positive values of PC1 and PC2, and grass seed abundance to positive values of PC3 (Fig. 2).

**Table 2.** Results of the Principal Components Analysis of the ten variables measured at the microhabitat scale. Main loadings for the three retained components are shown in bold.

|  | PC1 | PC2 | PC3 |
| --- | --- | --- | --- |
| Grasses | 0.334 | -0.138 | **0.648** |
| Dry standing grasses | 0.085 | 0.061 | **0.752** |
| Low shrubs | 0.502 | 0.339 | **-0.435** |
| Tall shrubs (< 1m) | **0.789** | -0.392 | 0.129 |
| Tall shrubs (> 1m) | **0.767** | -0.396 | 0.133 |
| Trees (< 1m) | 0.017 | **0.850** | 0.040 |
| Trees (> 1m) | 0.121 | **0.865** | -0.128 |
| No vegetation | **-0.872** | -0.162 | -0.176 |
| Bare ground | **-0.875** | -0.301 | -0.167 |
| Dense litter | **0.877** | 0.292 | 0.094 |
| Eigenvalue | 3.892 | 2.121 | 1.295 |
| % variability explained | 38.9 | 21.2 | 12.9 |
| % variability accumulated | 38.9 | 60.1 | 73.1 |

**Figure 2.**
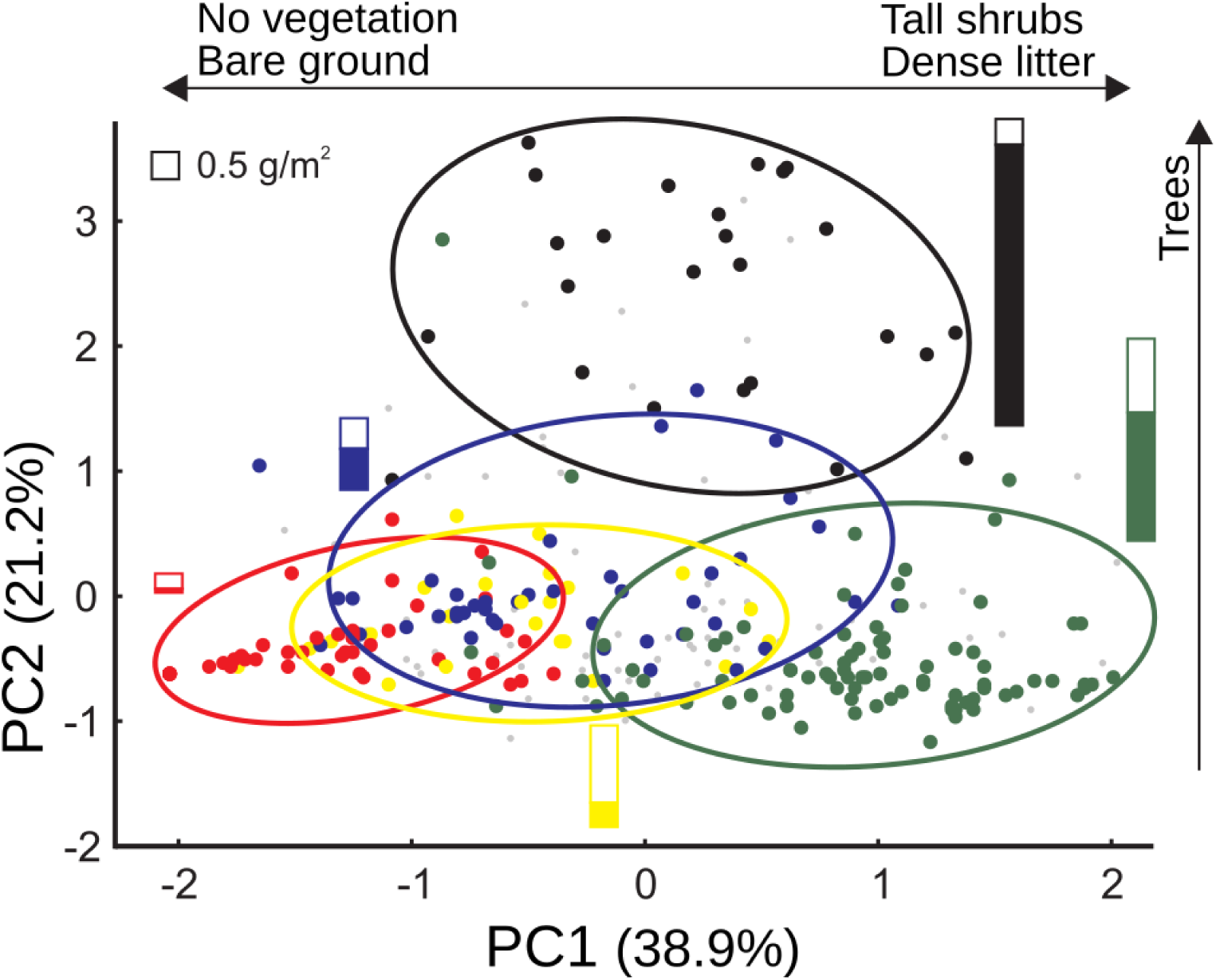
Position of the 300 seed-offer devices according to the first two components of a PCA on ten vegetation and soil variables measured at the microhabitat scale. Microhabitats are identified by their *a priori* categorization at field according to previous studies of the soil seed bank (Marone & Horno 1997, Marone & al. 2004): beneath tree canopy (black), beneath tall shrubs (green), beneath low shrubs (blue), beneath grasses (yellow) and bare soil (red). Smaller gray points were microhabitats with intermediate characteristics according to those criteria. Ellipsis include at least 90% cases in each category. Bars show the biomass of forb (full) and grass (open) seeds of each microhabitat category during the same winter of the field experiment (L. Marone, unpublished data). Relative sizes of bars look similar if number of seeds, instead of biomass, is represented (0.5 g/m^2^ ≈ 2000 seeds/m^2^).

The three grids were generally similar in their vegetation and soil characteristics at this scale though differed slightly on the distribution of their microhabitats over some PCA components: there were more microhabitats covered with low shrubs in V and with grasses in F (low and high values in PC3, respectively), and some more devices under trees in J than in V (PC2) (Kruskal-Wallis tests, *k* = 3, *n* = 300; PC1: *H* = 0.48, *P* = 0.78; PC2: *H =* 5.28, *P =* 0.07; PC3: *H =* 21.68, *P* < 0.01). Mean distances from devices to nearest tall trees showed higher differences among grids: devices in grid J were closer in average to a tree >3 m high (distance to nearest tree: 2.69 ± 2.27 m), F was intermediate (3.49 ± 3.05 m), and V devices were farther away in average (6.66 ± 4.45 m; K-W test: 48.69, *P* < 0.001). A similar though slightly stronger pattern was found for distances to only the tallest algarrobos (>4 m high; J: 5.60 ± 4.22 m; F: 8.63 ± 5.51 m; V: 13.18 ± 7.83 m; K-W test: 55.39; *P* < 0.001). Since this pattern matched observed levels of seed removal among grids, selection of characteristics associated with the devices were explored in analyses both pooling grids (i.e., analysis at the habitat extent) and within each grid.

### Foraging site selection based on microhabitat characteristics

There were used devices across all the multivariate space defined by the three PCA components on vegetation and soil characteristics at the microhabitat scale (Fig. 3). However, seed removal was not completely random: used devices in spring and summer (pairs N_11_) and non-used devices in spring (pairs N_00_) were more aggregated than expected by chance (Table 3).

**Table 3.** Analysis of segregation of used and non-used devices in the 3D-space generated by the first three components of a PCA on ten variables measured at microhabitat scale. The observed and expected number of pairs of nearest-neighbour points that were both used (N_11_) or non-used (N_00_) in each seasonal experiment are shown, together with the statistic (C) testing for global spatial segregation between classes of points against the null hypothesis of “random labeling”, and the statistic (Z, between brackets) testing the same hypothesis for each kind of pair. A significantly higher number of pairs observed than expected indicates positive spatial correlation (aggregation, in bold).

| | observed/expected | $C$ ($Z$) | $P$ |
| --- | --- | --- | --- |
| Winter |  | 0.271 | 0.873 |
| $N_{11}$ | 131/134.5 | (-0.510) | 0.610 |
| $N_{00}$ | 32/32.5 | (-0.083) | 0.934 |
| Spring |  | 12.747 | 0.002 |
| $N_{11}$ | **85/63.2** | **(3.380)** | **0.001** |
| $N_{00}$ | **103/87.2** | **(2.328)** | **0.020** |
| Summer |  | 5.895 | 0.052 |
| $N_{11}$ | **61/47.0** | **(2.339)** | **0.019** |
| $N_{00}$ | 119/109.0 | (1.464) | 0.143 |
| Autumn |  | 2.452 | 0.293 |
| $N_{11}$ | 114/123.9 | (-1.457) | 0.145 |
| $N_{00}$ | 38/37.9 | (0.012) | 0.991 |

**Figure 3.**
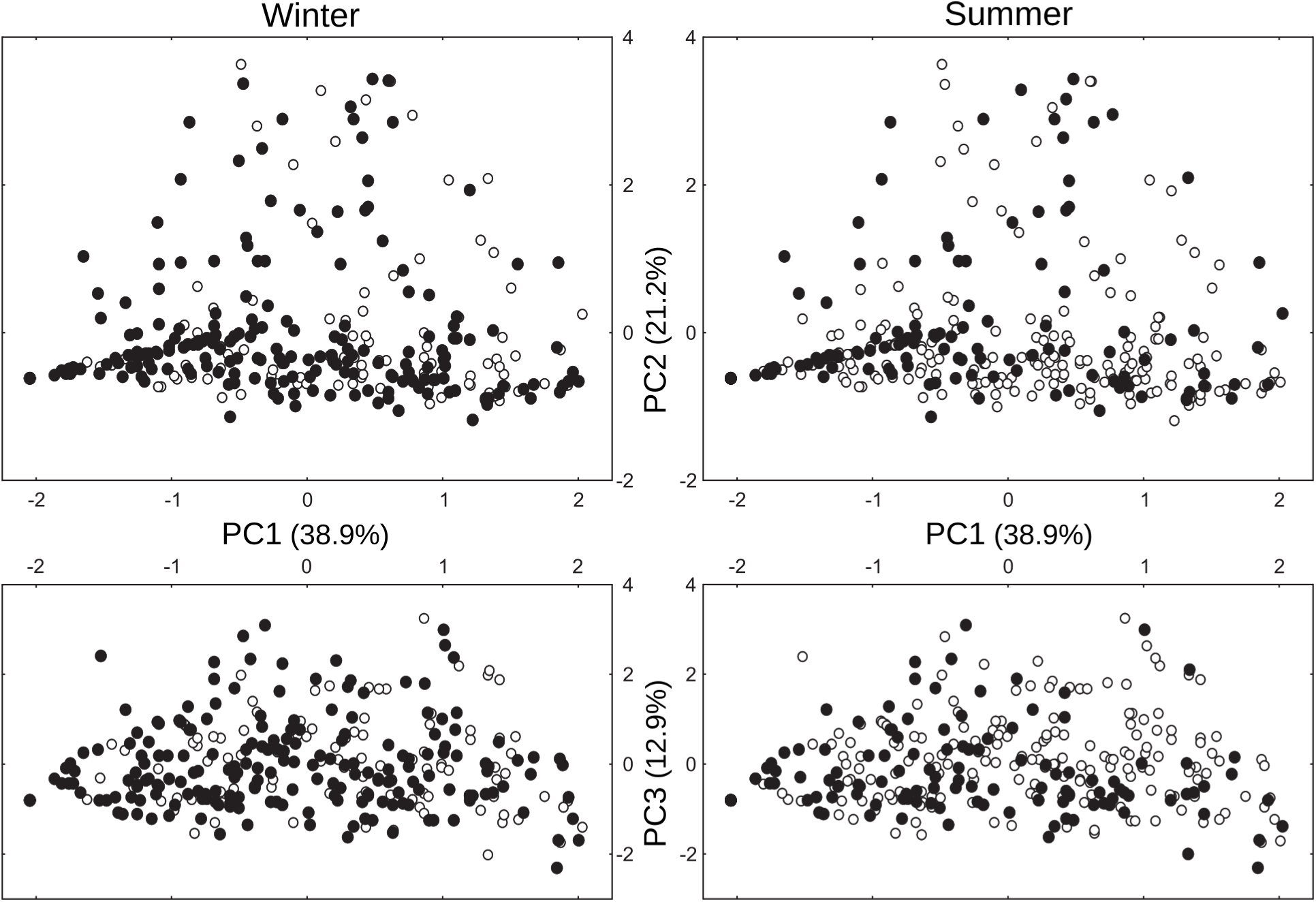
Distribution of used devices (filled circles) in the season with more (winter) and less (summer) seed removal in the multidimensional space of the three first components of a PCA on vegetation and soil characteristics at the microhabitat scale. Point patterns in autumn were similar to those in winter, and in spring to summer’s.

Unidimensional analysis along each PCA component also indicated that the probability of a seed being removed is not independent of the main characteristics at the microhabitat scale. In all seasons, there were some differences in one or more components between the mean value of explored microhabitats and the expected mean under the null model (Fig. 4). However, evidence of foraging selection depended on the null model assumed. In the analysis at the habitat extent, there was a preference for devices in microhabitats with less cover of shrubs and litter (PC1) in all seasons, in microhabitats with trees in spring and summer (PC2), and with less grasses (or more low shrubs) in summer (PC3). Variances of scores of used devices were no different from those expected by chance in any component or season, even when different means were detected (i.e., even when means differed, characteristics associated to used devices were as variable as those available). When the analysis controlled for the possible hierarchical selection at the grid level with a stratified null model, skewed use was still observed on PC1, favouring microsites without cover and litter in all season, but significant selection along PC2 practically disappeared and preference on PC3 for microhabitats with no grasses became stronger (statistically significative in spring, summer and autumn; Fig. 4).

**Figure 4.**
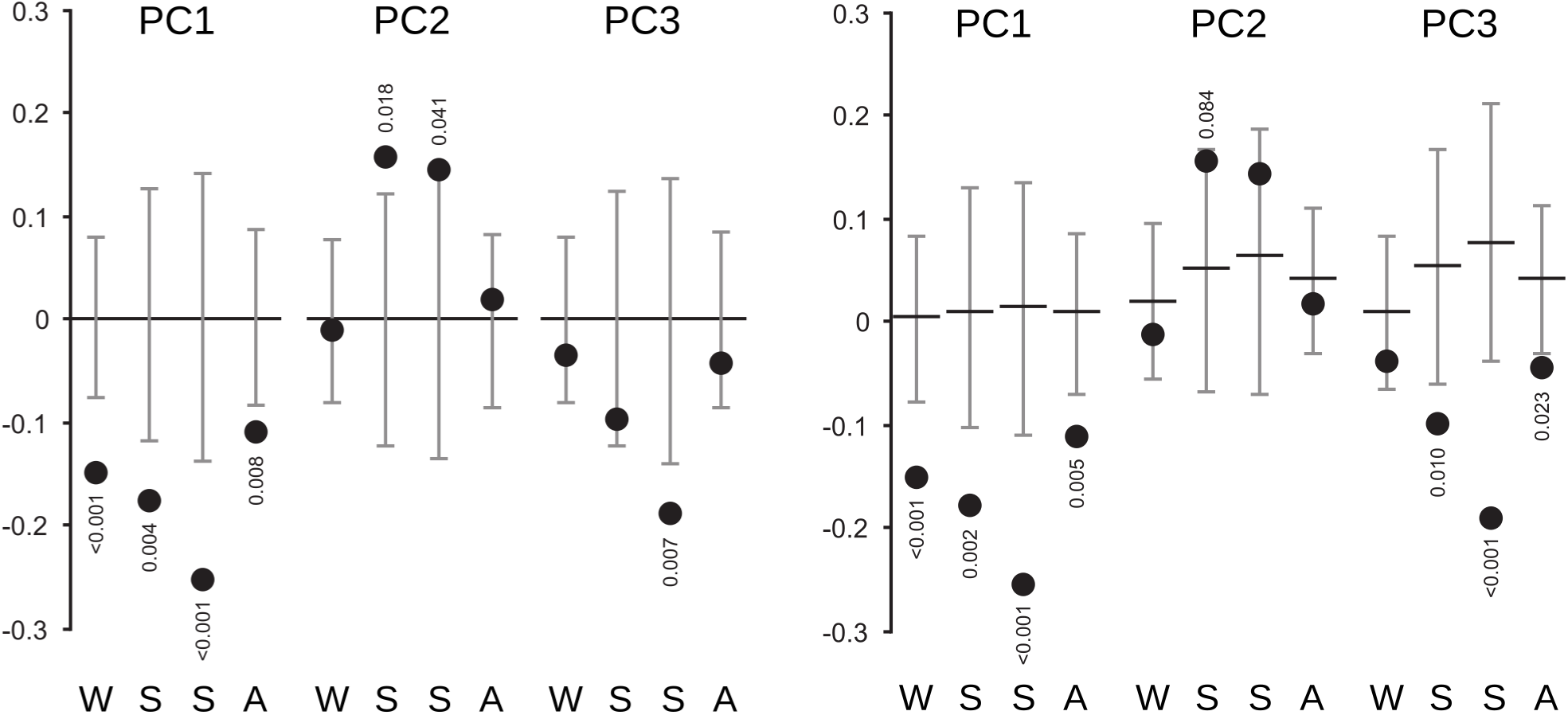
Mean scores of used devices in the first three components of the PCA (black circles) in each season (from winter [W] to autumn [A]), controlling (right panel) or not (left panel) for potential selective use of space at the grid scale. Black horizontal line is the expected value and whiskers show the 95% confidence interval under the null hypothesis of no selection. Pseudo P-values not shown when *P* > 0.1 (two-tailed test).

### Foraging site selection based on microhabitat position

In several seasonal trials there was a positive spatial association of seed removal at short distances (5–7.5 m). Neighbouring devices (i.e., separated by the first distance class) were both used more frequently in spring (*P =* 0.018) and autumn (*P =* 0.023) in grid F, and in spring (*P =* 0.006) and summer (*P =* 0.001) in grid V, than expected by chance (Join-counts analyses against CSR null model; Fig. 5). Similar patterns, though statistically non-significant, were observed for other seasonal trials in the same two grids (winter in F: *P =* 0.071; autumn in V: *P =* 0.103). No significant spatial pattern was detected in grid J. Correlogram slopes (i.e., how autocorrelation changes with distance) were always steeper in spring and summer than in autumn and winter (Fig. 5).

**Figure 5.**
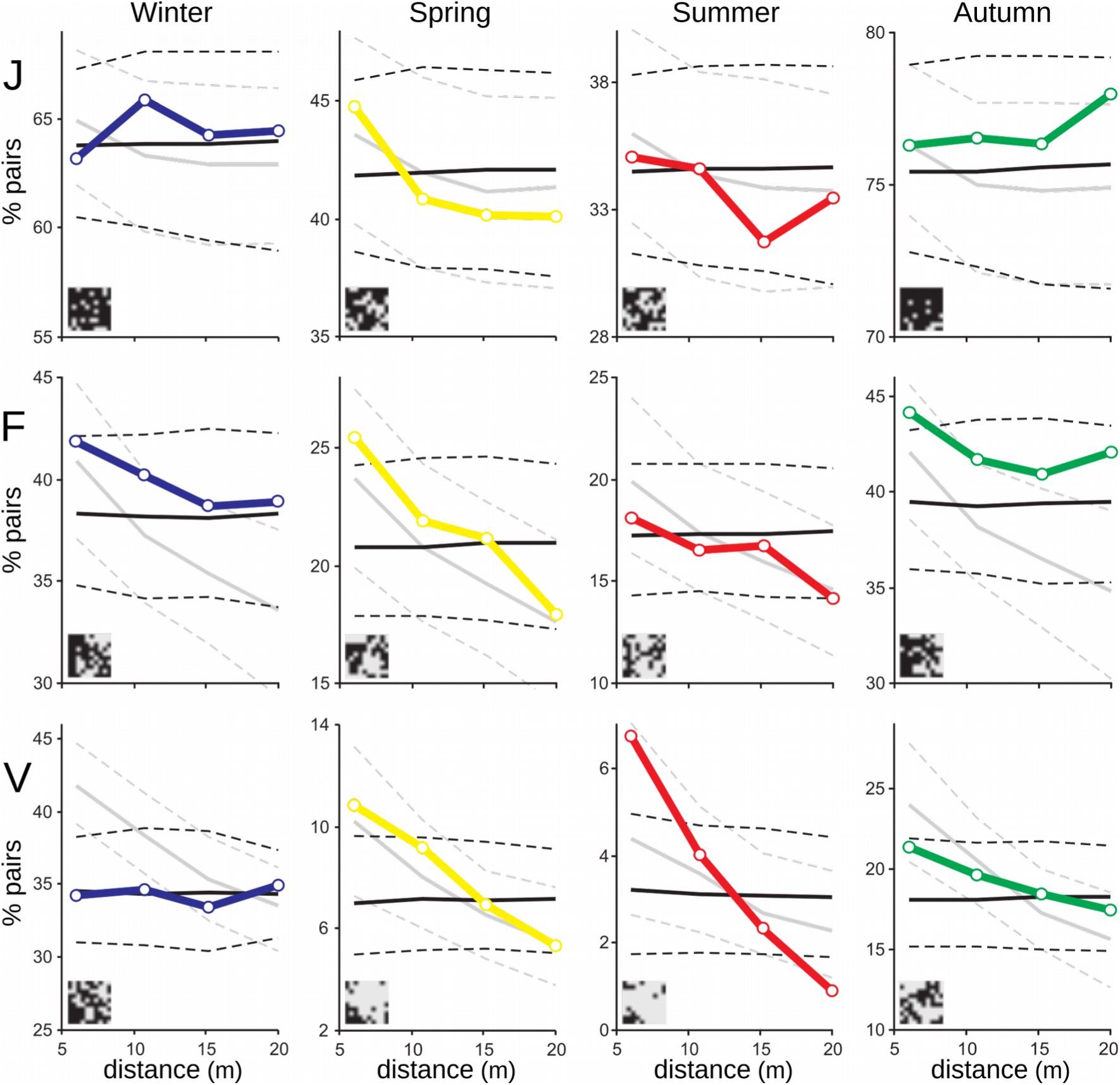
Percentage of used-used pairs of devices along four distance classes in each season in three 10×10 grids (J, F, V). Black lines show the expected percentage (continuous) and 2.5% and 97.5% percentiles (broken) of used-used pairs under a complete spatial randomness (CSR) model. Gray lines show the same three statistics under a spatially heterogeneous model that considers a negative influence of increasing distances to tall trees (see details in text). Actual patterns of seed removal are shown as insets in the bottom left corner of each subfigure, with used devices in black.

Spatial aggregation of seed removal was not a direct consequence of spatial autocorrelation of vegetation characteristics at the microhabitat scale (Fig. 6): (1) PC1, the component more consistently associated with seed removal (Fig. 4) was not spatially autocorrelated; (2) PC2 was positively autocorrelated in grid V at the first distance class but was not significantly relevant in explaining seed removal once the effect at the grid scale was controlled for; and (3) PC3 was positively autocorrelated in grid J and at the second distance class in F, both at which there was no evidence of autocorrelated seed removal (Fig. 5).

**Figure 6.**
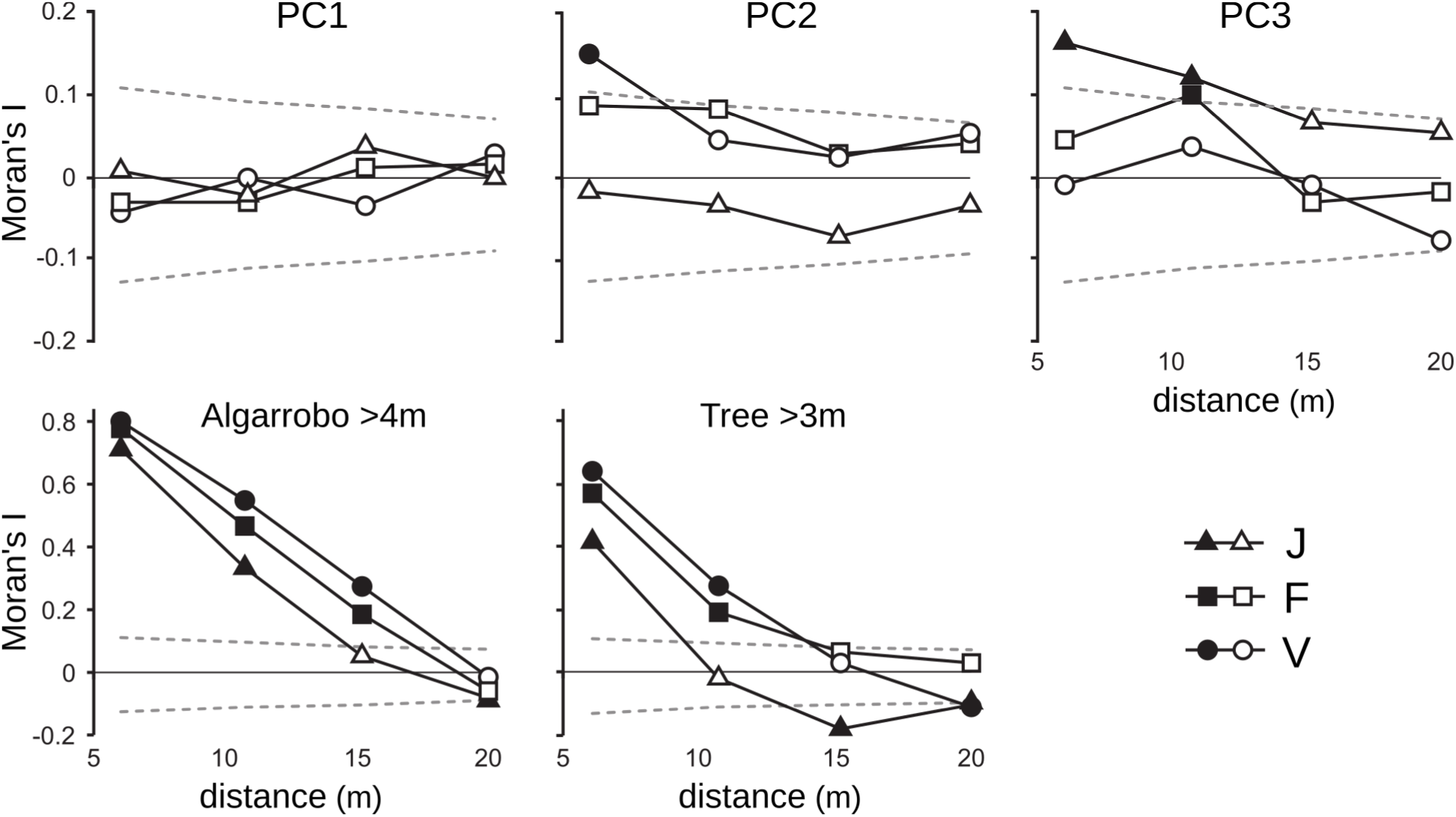
Correlograms of synthetic variables (after a PCA) with characteristics of the vegetation at microsite scale and of two variables of distance to tall trees in the three 10×10 grids (J, F, V). Broken lines are the 2.5% and 97.5% percentiles of the Moran’s I values expected by chance under a complete spatial randomness model (CSR); full symbols show values outside those point-wise intervals (i.e., *P* < 0.05).

In contrast, distance from devices to tall trees showed a strong positive autocorrelation (Fig. 6) as a consequence of many neighbouring devices sharing the same nearest tall tree (26.7–68.0% distances from a device to the closest tall algarrobo were >5 m). Correlograms describing autocorrelation of seed removal were more similar to those of a heterogeneous Poisson model with tall trees as main predictor (DTT model, Fig. 5). Observed correlograms in grids V and F were more similar to DTT model in spring and summer, particularly at short distances, while autumn-winter pattern were intermediate or more similar to CSR models, particularly at longer distances. In grid J both models had similar predictions as a consequence of the higher abundance and more uniform distribution of tall trees; no observed pattern would have statistically distinguished between the two null models.

## DISCUSSION

As a consequence of evolutionary or behavioral adaptation, foragers should be able to track spatial variations in food abundance when coupled to perceptible environmental cues (Bell 1990, Schmidt & Brown 1996, Fierer & Kotler 2000, but see Fawcett et al. 2013). We tested the hypothesis that exploration of foraging patches by postdispersal granivorous birds is guided by environmental features consistently associated with seed abundance in the soil of a desert. We offered scattered single seeds at field in seasonal trials to detect selective spatial patterns by foraging birds and analyze their association with vegetation and soil characteristics at the microhabitat scale. Our multivariate characterization of vegetation heterogeneity at microhabitat scale (1–2 m diameter) proved consistent among studies in this habitat (*algarrobales* at Ñacuñán, in the central Monte desert, Argentina; cf. Milesi et al. 2008) and matched the main vegetation features involved in criteria categorizing microhabitats for previous analyses of the soil seed bank (Marone & Horno 1997, Marone et al. 2004). Therefore, foraging spatial patterns at this scale can be interpreted according to our previous knowledge on seed bank abundance, composition and dynamics. We start by discussing the experimental field technique used and the consistent evidence it provided, then focus on our main hypotheses on selective patterns of seed removal by granivorous birds at the microhabitat scale, and finally discuss birds’ foraging patterns at bigger spatial scales, their seasonal variation and their potential causes and implications.

### Postdispersal granivory by birds as inferred from seed-offer field trials

From results of Mares & Rosenzweig (1978) onwards, several authors have assumed that desert birds rarely find, or are slow to detect, experimental seed dishes or trays (e.g., Abramsky 1983, Parmenter et al. 1984, Kerley 1991; see Marone et al. 2000b, Weisz et al. 2012). This would set a strong limit to short-term field experiments that avoid baiting or training birds, so it is not surprising that most seed-removal trials have exploited the “magic pudding effect” (Bedoya-Pérez et al. 2013) to, intentionally or not, emphasize the exploitation rather than the exploration component of foraging. Contrastingly, our experimental scattered seeds were readily found and removed by local birds during the first day of every seasonal trial, even when seed addition at the grid scale was nil (100 seeds in ~0.2 ha, or <1.5 g/ha). The relevance of small granivorous desert birds to exploit scarce, dispersed seed resources could be higher than previously considered, or at least it should not be assumed poor by extrapolating observations from other regions.

The identification or control of selective access by different granivorous taxa is a typical problem in seed bait studies (Weisz et al. 2012), usually involving physical barriers that do not always prove successful (Christianini & Galetti 2007, Brown et al. 2016, *pers. obs.*). We tested and used a day–night criterion calibrated by recording foot tracks over fine soil around ant-proof seed-offer devices. In this environment, single-seed offer proved a very simple, quick, scalable and economic solution for studies with higher resolution or spatial extent, and facilitates the assumption that experimental seed removal reveals the usual use of foraging space by granivorous birds. Radiotelemetry (e.g., Macias-Duarte & Panjabi 2013) or videographic techniques (e.g., Mokotjomela et al. 2014, Brown et al. 2016) may be more precise or complete alternatives (see also Bedoya-Pérez et al. 2013), though they are more expensive, time-consuming and limited to a minimum scale of resolution or a lower number of focal places.

Seasonal variation in seed removal by birds proved consistent within and across studies, adding another evidence in support of this technique to estimate postdispersal granivory. Seed removal was always highest in autumn and winter and decreased in spring to a minimum in summer. This seasonal pattern, repeated in each grid of our field study, was also evident after direct observations of small granivorous birds foraging on the ground (Milesi et al. 2008) and by measuring seed biomass removed from 25–50 *ad libitum* trays (Lopez de Casenave et al. 1998; see figure 3.21 in Milesi 2006 for a graphical comparison of these results). Reported local changes in abundance and diet of these small granivorous birds also match. The granivorous guild changes along the year with the arrival of some migrant birds during autumn and winter, particularly the abundant southern subspecies *Zonotrichia capensis australis* (Sagario et al. 2014). Though the arrival of *Poospiza ornata* for the breeding season (Cueto et al. 2011) partially compensates the departure of *Z. c. australis* in spring, all postdispersal granivorous birds reduce their seed intake in spring and summer to include insects and fruit in their breeding diets (Lopez de Casenave 2008, Milesi et al. 2008).

Spatial patterns of seed removal may be wrongly inferred if seed quantity or quality in the soil around the experimental device correlate negatively with the probability of removal of the focal seed (safety-in-numbers or dilution effect): patch use would appear higher in poor microhabitats even when birds were spatially non-selective. However, this interpretation does not seem probable for a number of reasons: (1) these experimental seeds are taxonomically similar to some naturally available but preferred and abundantly removed when offered *ad libitum* (e.g., Lopez de Casenave et al. 1998), (2) removal of single seeds did not differ from removal of groups of ten seeds in a pilot study, suggesting that birds completely remove groups of experimental seeds when they are easily available in an accessible artificial patch, and (3) we never found bird footprints around a device with the seed unremoved in our additional field trial (see *Methods*). Notwithstanding, inferences on use of foraging space may include an experimental artifact if, in spite of precautions (device surface covered in natural soil, single seeds offered in short and isolated trials to reduce reward and minimize learning), seed devices were still particularly attractive for birds. Even if true, this should still reveal the spatial preferences of these birds when exploring their habitat for suitable foraging patches.

### Bird exploration and microhabitat characteristics

Postdispersal granivorous birds were less selective in their use of foraging space than expected if patch appearance (i.e., visual, remotely-perceivable environmental surrogates that provide reliable information on relative patch profitability) were providing them information to avoid poor or unprofitable patches. Variability in main microhabitat features was similar around used and available devices, suggesting that no kind of microhabitat is safe from bird exploration. Still, seed removal was not spatially random. Birds repeated the use of experimental devices at particular positions in consecutive days of each trial and along the seasonal trials in spite of intra-annual variations in seed abundance and distribution and in bird abundance, guild composition and behavior. This suggests that some permanent features associated to each individual foraging patch may still have a (non-restrictive) influence on its probability of exploration. Strikingly, the only consistent selective pattern, analyzed at both habitat and grid extents, was against previous expectations based on bottom-up behavioral effects. Microhabitat exploration in all seasons was skewed towards those with less perennial vegetation and litter cover, and in most cases with less grasses, all characteristics associated with fewer seeds in the soil bank. This selective use of space by birds at microhabitat scale allows an interpretation closer to a top-down spatial effect, a cause (and not a consequence) of the seed bank dynamics. Marone et al. (2004) reported grass seed abundance in the soil becoming more homogeneous among microhabitats with time after primary dispersal, and Marone et al. (2008) found that granivory may account for up to a 50% reduction of preferred, naturally accumulated grass seeds in open microhabitats.

During summer and autumn, birds may be targeting recently produced seeds that preferentially enter the seed bank through open spaces during primary dispersal (Marone et al. 1998a). Under this scenario, a more appropriate bottom-up model for microhabitat use by granivorous birds should focus on seed renewal rates (Price & Joyner 1997, Ben-Natan et al. 2004) rather than on the accumulated soil seed bank (following the general approach of an ideal free distribution model), or involve spatial autocorrelation or context at bigger scales to examine revisiting strategies such as cropping or traplining (Bell 1990). Still, foraging patterns from winter to early summer (when seed renewal rates are low because primary dispersal of grass seeds have already ended and most seeds were already trapped in litter or small depressions in the soil during secondary dispersal by wind and water; Marone et al. 1998a), do not seem to match predictions based on neither new nor accumulated seeds on the soil. Instead, birds may be guiding their search for seeds towards bare soil patches because of their detectability or accessibility (Whittingham & Markland 2002, Jones et al. 2006, Baker et al. 2009). Surface seeds on bare soil may be more easily detected or extracted by most local bird species than those trapped within dense litter under woody cover (Cueto et al. 2013).

In summary, according to their patterns of single-seed removal, granivorous birds in the algarrobal of the central Monte desert seem to explore every kind of microhabitat, ignoring visual information embedded in vegetation structure and soil cover to optimize their exploration for profitable foraging patches (behaving as myopic foragers *sensu* Mitchell 1989). The conclusion that they are unable to perceive bold characteristics of the vegetation is not straightforward, nor that their cognitive abilities do not allow them to recognize the distribution of patch qualities or develop a patch-searching rule based on them. Postdispersal seed foraging in a desert, i.e., one with high spatial variability and low encounter stochasticity (abundant small-sized prey) in a temporally changing environment, should favour the evolutionary value of learning over that of fixed strategies and parameters for the improvement of patch assessment rules (Eliassen et al. 2009). Then, those same evolved behavioural mechanisms to cope with this spatio-temporal variation (e.g., sample every patch so to increase survival when under extreme conditions) may not allow birds to satisfy predictions of optimality under a locally fixed, more beneficial set of features (the behavioral gambit *sensu* Fawcett et al. 2013). A fixed or narrowly defined patch selection template may not be convenient (enough) in a variable and dynamic system (Rotenberry & Wiens 1998) where a less-informed strategy may be ‘good enough’ (Olsson & Brown 2010). It is also probable that the foraging scale of these birds is much smaller than the spatial scale at which perennial vegetation provides useful information (see Kohlmann and Risenhoover 1998, Fierer & Kotler 2000, Klaassen et al. 2006a, 2006b). Even when in the algarrobal the general association between seed bank and vegetation is strong and persistent at microhabitat scale (decimeters to meters; Marone & Horno 1997, Marone et al. 1998a, 2004, Milesi & Marone 2015, Andrade 2016), patches at the relevant scale for the birds may be more variable than we acknowledge, as a consequence of accumulation gradients, micro-topography, secondary dispersion and continuous consumption. In this scenario, general *static* information embedded in woody vegetation would lose value to guide exploration at the relevant scale for the birds: (nano)patch quality would became less predictable, with instant sampling required anyway to search for and evaluate profitable patches (Vásquez et al. 2006, Stephens 2007, Eliassen et al. 2007, Reynolds 2012). Additionally, the penalty of unfruitful exploration of poor patches may be counterbalanced by the ability to detect scattered rich patches with structural characteristics associated with low-quality areas, such as ephemeral patches of recently-fallen grass seeds or scattered depressions accumulating seeds in exposed areas (an usual feature in Ñacuñán and other deserts: Price and Reichman 1987, Marone et al. 2004). Milesi & Marone (2015) tested for other important factors (accessibility to foraging patches, foraging efficiency, thermoregulation or predation risk) that may affect patch selection by birds at microhabitat scale, and although seed abundance was the main predictor, they also found sparrows frequently exploring the poor/variable patch, a behaviour that can be expected when not under immediate risk of starvation (Dall & Johnstone 2002) or as an attempt to avoid learning predators by being unpredictable in space (Lima 2002, Mitchell & Lima 2002).

### Bigger scales: influence of context on foraging patch exploration

Bigger spatial scales proved more relevant to describe heterogeneity in bird foraging in this habitat. Seed removal differed consistently among three grids (~0.2 ha) that were haphazardly established in the algarrobal, even when they proved similar according to main characteristics of the vegetation at the microhabitat scale. In the most used grid (“J”) every microhabitat appeared “suitable enough” (only 4 out of 100 devices were never used) while in the other two (“F” and “V”) birds behaved more selectively. In fact, differences in the amount of seeds removed from each of our grids were even higher than between seasons, which was a strikingly strong and repeatable pattern itself (illustrating an additional challenge for attempts at global explanations of granivory rates derived from one-shot seed-offer experiments; e.g., Folgarait et al. 1998, Peco et al. 2014). Moreover, seed removal in autumn-winter showed a random pattern at longer intra-grid distances with a tendency for aggregation at small distances, but in spring-summer it concentrated in areas of a relatively constant size, leaving the rest almost without exploration. In consequence, inferences on the selective use of microhabitats depend on how birds’ use of space at bigger scales is assumed. Since the first PCA component was not spatially autocorrelated at ≥5 m, inferences are not modified when controlling for spatial dependencies, but selective patterns identified along the other PCA components may follow from selective use at bigger scales. For example, the preference for tree microhabitats in spring and summer under the simplest analysis (at the habitat extent, i.e., pooling grids) disappeared when controlling for the differential use of each grid (with a stratified analysis or taking any of the grids as extent). It is well established nowadays that most ecological patterns depend on scale, including habitat selection and foraging patterns resulting from the combination of degrees of preference at different scales (Jones 2001, Morrison et al. 2006, Lopez de Casenave et al. 2007, Mayor et al. 2009). Although the evaluation of spatial dependences at bigger spatial scales than microhabitat was not our main goal, the influence of the scale of analysis on inferences suggests a selective use of foraging patches based not only on the characteristics of a device but also in its relative position among other devices or in relation to other structures.

Most notable among those influences was the negative association between seed removal by birds and distance to tall trees, both as a selective pattern among grid areas and within grids with enough variation for this variable. A simple heterogeneous Poisson model in which probability of seed removal was negatively related to distance to tall trees (assuming no further second-order interactions) successfully predicted a similar change in autocorrelation with distance in the two grids where it was observed and no autocorrelation in the third. A similar selective use of space around tall trees was also found by following foraging birds at field both for predispersive (directly from the spikes) and postdispersive (from the soil) seed consumption (Milesi et al. 2008) and it is not unusual elsewhere (e.g., Pulliam & Mills 1977). This preferential exploration of areas close to tall trees (but not exclusively under trees) does not seem related to seed abundance. Though we lack data on heterogeneity of seed abundance at this scale (e.g., tens of metres) or at different distances from trees within the algarrobal, previous local studies found no correlation between abundances of grass seeds and granivorous birds measured at a slightly bigger scale (200×100 m; Lopez de Casenave 2001), and average seed abundance was not very different even between close habitats differing markedly in tree cover (Marone et al. 1997, 2004). Birds may have preferred shaded patches beneath and around trees during summer for thermorregulation. Lopez de Casenave et al. (1998) also found birds being able to remove seeds at similar rates from *ad libitum* trays in both open patches and beneath trees in this habitat during winter but favoring seed removal under cover in summer. We evaluated this idea in an additional summer trial in grids J and V, with the same experimental design but tracking seed removal throughout the day (Milesi 2006). We observed (1) higher seed removal closer to tall trees in the heterogeneous grid (V) and no spatial pattern in the homogeneous one (J); (2) most seed removal occurred during the morning and afternoon and it was not related to shade; and (3) during midday hours, when shade becomes critical, seed removal was nil, even under cover (detailed results in Milesi 2006). Milesi & Marone (2015) also showed in controlled trials with single birds in field aviaries that extreme temperature or direct sunlight may pose a limit on foraging but can not be invoked as the single main reason for spatially selective foraging patterns.

Reproduction and predation are more probable causes of these spatial patterns. Autocorrelation of seed removal coupled with higher probabilities of exploration closer to tall trees is to be expected if granivorous birds have reproductive territories or home ranges set around them. Finches and sparrows may be using these tall trees as important posts for antipredatory and territorial vigilance. The (stronger) association with trees in the reproductive season cannot be explained by a central foraging model around nest sites since most nests of these granivores are not built on tall algarrobo trees but on shrubs, on the ground or in smaller *Geoffrea decorticans* trees (Mezquida 2003, 2004). Still, these small birds are territorial, particularly during spring and summer, when tall trees are preferred singing perches (Sagario 2011, Sagario & Cueto 2014a, Zarco 2016). Seasonal variations in degree of aggregation around trees may also associate with changes in guild composition, movement patterns, territoriality and territory size, e.g., multispecies flocks in autumn-winter with high relative abundance of the migratory subspecies *Zonotrichia capensis australis* (Sagario & Cueto 2014b, Sagario et al. 2014, Zarco & Cueto 2017). Affinity for tall trees may also be related to real or perceived predation risk, as interpreted in classic studies of foraging granivorous birds (e.g., Pulliam & Mills 1977, Schneider 1984, Lima et al. 1987, Schluter 1988, Watts 1991). Most local birds flee to the closest tall trees when disturbed within the algarrobal (*pers. obs.*) as frequently observed in other habitats with multi-layered vegetation structure. Foraging farther from tall trees may be associated with a higher risk of not reaching protective cover quick enough if attacked by a predator. This will trade-off against foraging related tasks, resulting in higher costs in terms of fitness (Brown & Kotler 2004, Verdolin 2006, Cresswell 2008). Further, seasonal variation in degree of aggregation around trees may relate to varying perception, evaluation or actual risk of predation (Watts 1991, Suhonen 1993, Lima & Bednekoff 1999). There was some support for the predation risk hypothesis in results of foraging selection trials within field aviaries by Milesi & Marone (2015).

Irrespective of which is the main cause, tall trees seem to determine a first level of selection that defines explorable space, and then microhabitat structure exerts an influence on which patches are effectively exploited (or more frequently explored) among the accessible ones. This is not an unusual conclusion when studying small ground-foraging birds (e.g., Schluter 1988, Repasky & Schluter 1994, Watts 1996, Walther & Gosler 2001). In our case, openness (no woody cover), probably as a cause of better accessibility at ground level or to facilitate predator detection, seems more important at this smaller scale than vegetation features correlated with soil seed abundance. Moreover, this two-step explanation matches the two movement modes of these birds: flying between perches and from perches to the ground, and then walking on the ground while foraging. Short term seed-removal trials may not be long enough to reveal the existence of predation-free areas for seeds, but may still estimate their relative probability of being consumed according to the intensity of exploration of foraging patches by birds within their activity areas. Even when territories of granivorous birds in the algarrobal tend to be contiguous (Sagario 2011), a selective use of space as hypothesised may cause a scenario of heterogeneous risk of predation for seeds (particularly for seed grasses preferred by birds), such as scarcely used patches far from tall trees. Then, heterogeneous removal of seeds in a seed-limited environment (this habitat: Andrade 2016) may cascade to the spatial distribution of plant populations. Expected foraging patterns under this multiscale hypothesis associated with birds’ two-step movements should ideally be tested at bigger spatial scales, including higher degrees of environmental heterogeneity and more diverse distances to tall trees.

## ACKNOWLEDGEMENTS

We thank L. Marone for partnership and valuable guidance at several stages. We acknowledge institutional and partial financial support from Aves Argentinas, CONICET, Universidad de Buenos Aires, IADIZA (CONICET-UNCuyo-Mendoza), and ANPCyT, all from Argentina. This is contribution number 103 of the Desert Community Ecology Research Team (Ecodes) of IADIZA (CONICET) and FCEN (Universidad de Buenos Aires).

